# Epigenetic control of multiple genes with a single lentiviral vector encoding transcriptional repressors fused to compact zinc finger arrays

**DOI:** 10.1101/2024.01.17.576049

**Authors:** Davide Monteferrario, Marion David, Satish K. Tadi, Yuanyue Zhou, Irène Marchetti, Caroline Jeanneau, Gaëlle Saviane, Coralie F. Dupont, Angélique E. Martelli, Lynn Truong, Jason Eshleman, Colman Ng, Marshall Huston, Gregory D. Davis, Jason D. Fontenot, Andreas Reik, Maurus de la Rosa, David Fenard

**Author notes:** **Correspondence :** David Fenard, PhD, Sangamo Therapeutics France, Les Cardoulines HT1, Allée de la Nertière, 06560 Valbonne France. **Email :** or.

## Abstract

Gene silencing without gene editing holds great potential for the development of safe therapeutic applications. Here, we describe a novel strategy to concomitantly repress multiple genes using zinc finger proteins fused to Krüppel-Associated Box repression domains (ZF-Rs). This was achieved via the optimization of a lentiviral system tailored for the delivery of ZF-Rs in hematopoietic cells. We showed that an optimal design of the lentiviral backbone is crucial to multiplex up to three ZF-Rs or two ZF-Rs and a chimeric antigen receptor. ZF-R expression had no impact on the integrity and functionality of transduced cells. Furthermore, gene repression in ZF-R-expressing T cells was highly efficient *in vitro* and *in vivo* during the entire monitoring period (up to ten weeks), and it was accompanied by epigenetic remodeling events. Finally, we described an approach to improve ZF-R specificity to illustrate the path towards the generation of ZF-Rs with a safe clinical profile. In conclusion, we successfully developed an epigenetic-based cell engineering approach for concomitant modulation of multiple gene expressions that bypass the risks associated with DNA editing.

## INTRODUCTION

Epigenetic modifications play a critical role in the regulation of gene expression. They are heritable and reversible changes that affect gene expression without altering the DNA sequence. In mammalian cells, Krüppel Associated Box (KRAB)-containing zinc finger (ZF) proteins (KRAB-ZFPs) represent a family of more than four hundred DNA-binding transcriptional repressors.^1^ These domains are crucial for the epigenetic regulation of transposable elements and mammalian evolution.^2, 3^ The interaction of KRAB-ZFPs with their genomic target locus allows the recruitment of critical co-factors like KAP1/TRIM28,^4^ and consequently, the epigenetic transcriptional silencing of the target gene through various mechanisms, including DNA methylation and histone modifications. The latter involves the addition or removal of chemical groups to the histone tails (*e.g.* methylation, deacetylation), which can influence the accessibility of DNA to the transcriptional machinery.^5^

To specifically target a KRAB domain to a locus of interest in the genome, different DNA-binding modalities have been developed.^6^ They can be derived from the CRISPR/Cas9 system, in which the dead Cas9 protein is fused to a KRAB domain (dCas9-KRAB) and requires co-expression of a guide RNA capable of interacting specifically with the target locus.^7, 8^ The DNA binding protein can also consist of a transcription activator-like effector protein fused to a KRAB domain (TALE-KRAB).^9, 10^ And finally, the DNA-binding domain can be derived from an engineered ZFP, containing multiple zinc-fingers in a single protein, a technology developed nearly thirty years ago.^11–13^ Among all these DNA-interacting technologies, the last generation of engineered ZFP has the advantage i) to target virtually any DNA sequence in the genome;^14^ ii) to be very compact, iii) to be an all-in-one technology, harboring DNA binding capacity and KRAB domain on the same protein and iv) to be of human and not bacterial origin, decreasing the chance of a pre-existing or induced immune response for therapeutic applications.^15–17^ Engineered ZFP-KRAB repressors (ZF-Repressors, ZF-Rs), allow the precise repression of any target gene.^18–21^ ZF-Rs have been developed in AAV gene therapy approaches targeting the central nervous system for the treatment of tauopathies^20^ or huntington’s disease.^21^ While the AAV delivery system is adapted for non-dividing cells like neurons, this episomal delivery of the ZF-Rs is not suitable for constitutive expression in highly dividing hematopoietic cells. For the latter, the use of third-generation HIV-1-derived lentiviral vectors (LVs) pseudotyped with the vesicular stomatitis virus envelope glycoprotein (VSV-G) is more adapted and extensively employed in *ex vivo* hematopoietic cell therapies.^22, 23^ Furthermore, the lentiviral packaging capacity extends the multiplexing capacity of ZF-Rs compared to AAV. In this study, we optimized a lentiviral backbone design for an efficient multiplexing of up to three ZF-R sequences, allowing stable epigenetic control of multiple genes in T cells or regulatory T cells (Tregs) after a single lentiviral transduction event.

## RESULTS

### ZF-Repressor design and characterization

To evaluate the ZF-R technology efficiency, multiplexed in a single lentiviral particle, four human genes were selected for their potential therapeutic application. These genes are the beta 2-Microglobulin (*B2M*) and the class II major histocompatibility complex transactivator (*CIITA*), which regulate cell surface expression of the major histocompatibility complex class I (MHC-I) and class II (MHC-II) respectively, and are often knocked out in allogeneic T-cell products to prevent rejection or GvHD ;^24, 25^ the *CD3zeta* (*CD3Z*) gene, a component of the T cell receptor (TCR)/CD3 complex, essential for its expression and signaling;^26, 27^ and the *CD5* gene, a target repressed in cell therapy expressing an anti-CD5 chimeric antigen receptor (CAR) to prevent fratricide.^28^ For each target gene, approximately 200 research-grade ZFPs targeting the flanking regions of the transcriptional start site (TSS) were designed and linked to the human KOX1 KRAB repressor domain. The repression efficiency of these ZF-Rs was evaluated in human T cells two days after ZF-R mRNA electroporation by quantifying target gene mRNA expression using reverse transcriptase-quantitative polymerase chain reaction (RT-qPCR). As shown in Figure 1 and Figure S1, highly efficient ZF-Rs for all four target genes were identified in these screens. The targeted *B2M* locus showed that only a small set of ZF-Rs targeting the exon 1 region was highly efficient (Figure 1A), highlighting the importance of designing an extensive library of ZFPs to target efficiently a large portion of the TSS region. The ZF-Rs targeting CD5 (Figure 1B), CD3zeta and CIITA (Figure S1) exhibited a potent repression across a wider genomic region neighboring the TSS. In later screens, alternative KRAB domains^7^ were tested and several of them (*e.g.* human ZIM3, ZNF324) harbored an equal or improved repression activity compared to KOX1 (data not shown). Finally, in line with the transient nature of mRNA transfection experiments, the transcript expression levels of target genes returned to basal levels within few days, confirming that stable ZF-R expression in the rapidly dividing T-cells is required for a durable gene silencing.

**Figure 1.**
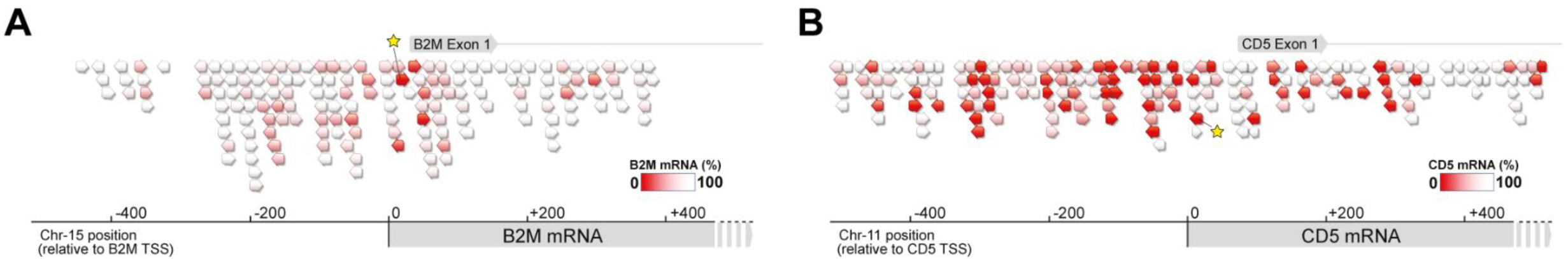
Screening of functional B2M and CD5 ZF-Repressors in primary T cells. Schematic of the *B2M* (A) and *CD5* (B) target genes with ZF-R candidates binding near the transcription start site (TSS). Triangles represent ZF-R binding locations on the target gene and the red color intensity correlates with repression efficiency evaluated by RT-qPCR two days after ZF-R mRNA transfection in T cells. The triangle orientation indicates the binding orientation of ZF-Rs. The selected lead ZF-Rs are highlighted by a yellow star.

### Bidirectional lentiviral vector design for efficient ZF-R multiplexing

To stably integrate ZF-R sequences in the genome, we set out to generate a lentiviral delivery system. We initially aimed for an efficient, easily interchangeable LV design allowing functional co-expression of up to two ZF-Rs, and a truncated version of the nerve growth factor receptor (dNGFR) to monitor and enrich transduced cells. Strategies to multiplex transgenes can be based on the use of picoviral self-cleaving 2A peptides.^29–31^ Despite some functionality alone in a bicistronic setting, 2A peptides have shown reduced efficiency when combined within a multicistronic cassette, resulting in a gradual loss of transgene expression towards the end of the 2A-linked peptide chain.^30, 32^ To mitigate this potential limitation, we generated a bidirectional LV architecture, a typical approach in LV backbone design.^33, 34^ It consists of an antisense cassette expressing the dNGFR protein, and a sense cassette accommodating codon-diversified ZF-Rs, targeting *B2M* and *CD5* genes either in simplex or duplex format connected by a self-cleaving *thosea asigna* virus 2A (T2A) sequence (Figure 2A). Diverse KRAB domains, such as the transcriptional repressor domain of KOX1, ZNF324 and Zim3,^7^ were used for ZF-R design to avoid repeated sequences that could be deleterious for LV production and infectivity upon multiplexing (Figure S2). Furthermore, the human phosphoglycerate kinase (PGK) and elongation factor 1 alpha (EF1a) promoters have been selected to form bidirectional transcriptional units as described previously.^34^ To identify the desirable lentivector design ensuring robust repression of the indicated ZF-R targets, we first compared two different LV architectures, each harboring the described transcriptional units in inverted orientation with respect to each other (pLV.PGK-EF1a and pLV.EF1a-PGK). Following T cell transduction, both LV architectures resulted in a comparable genomic integration capacity (*i.e.* VCN per cell) regardless of the cargo (Figure 2B, left panel). Consistently, dNGFR positive cells were detected nearly at the same frequency, indicating that the orientation of the transcriptional units did not impact the transduction efficiency (Figure 2B, right panel). ZF-R mediated gene repression was next assessed by flow cytometry on dNGFR-enriched cells at D10 post-transduction. Monocistronic sense cassettes resulted in high percentages (>90%) of CD5(-) or MHC-I(-) cells, and no significant differences were observed when comparing the two LV architectures (Figure 2C). In contrast, bicistronic sense cassettes were less efficient in co-repressing MHC-1 and CD5, with the EF1-PGK bidirectional promoter resulting in a dramatically reduced percentage of CD5(-)/MHC-I(-) cells as compared with its inverted counterpart (19.4% vs 62.6%, respectively). These findings were further corroborated by RT-qPCR, revealing a significantly more pronounced decrease of *CD5* and *B2M* mRNA transcripts when ZF-R expression was driven by the EF1a promoter, rather than PGK (Figure S3). To further optimize this LV design, the PGK promoter was replaced with a previously reported shorter version, namely PGK200,^35^ with the intent to reduce cargo size and improve infectivity (Figure 2A and Figure S2). This PGK200-EF1a backbone improved LV infectivity as shown with the slight increase in VCN per cell as well as the higher percentages of dNGFR positive cells compared with PGK-EF1a backbone (Figures 2B). In addition, constructs harboring the bicistronic sense cassette showed a striking increase in the frequency of CD5(-)/MHC-I(-) cells (>80%), suggesting that the indicated modification did not only improve transduction (Figure 2B), but also repression efficiency and with nearly no impact on dNGFR expression levels (Figure S4). To repress three different genes simultaneously with a single LV particle, the dNGFR sequence was replaced with a ZF-R sequence targeting the CD3zeta promoter (Figure 2D and Figure S2). In line with previous studies showing that CD3z is required for the cell surface expression of the TCR/CD3 complex,^26, 27^ this triple ZF-R construct resulted in the concomitant downregulation of MHC-I, CD5 and the TCR/CD3 complex (Figure 2D and figure S5). Altogether, these data defined the pLV.PGK200-EF1a architecture as an LV backbone of choice for efficient ZF-Rs multiplexing.

**Figure 2.**
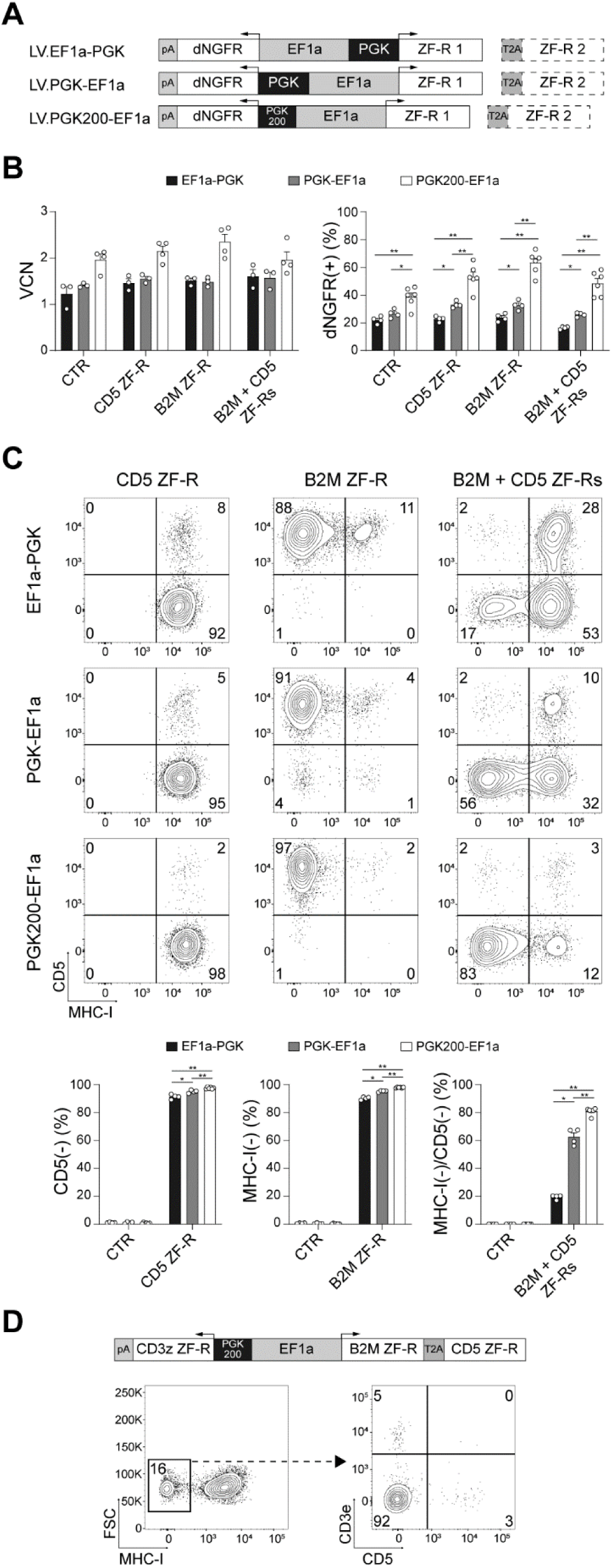
Lentiviral vector backbone design for efficient ZF-R multiplexing. A) Schematic representations of bidirectional LV backbones harboring two promoters, either EF1a-PGK, PGK-EF1a or PGK200-EF1a to co-express the dNGFR marker and one or two ZF-Rs. The different bidirectional LV backbones express either no ZF-R (CTRL), a single ZF-R (CD5 ZF-R or B2M ZF-R) or a combination of CD5+B2M ZF-Rs. B) Transduction efficiency of the indicated LV vectors was assessed by measuring the vector copy number per cell (VCN, left panel) and the percentage of dNGFR+ cells by flow cytometry (right panel) at D4 post-transduction. Data are represented as mean ± SEM. White circles indicate data obtained from each T cell donor (n=4-6). C) CD5 and MHC-I repression in dNGFR-enriched T cells transduced with the indicated LV vector was assessed by flow cytometry at D10 post-transduction. Data are represented as the mean ± SEM (n=4-6 donors). D) Representative plots showing concomitant MHC-I, CD3 and CD5 repression in T cells transduced with the triple ZF-R lentivirus at D12 post-transduction. The Mann-Whitney test was performed and only statistically significant differences were indicated: *\*P* ≤ 0.05*; **P* ≤ 0.01.

### ZF-R mediated silencing of the *B2M* gene is associated with repressive epigenetic mark

To interrogate the chromatin-modifying activity of the B2M ZF-R, we compared the epigenetic status of the *B2M* locus in T cells transduced with a bidirectional LV vector expressing either B2M-ZF-R or no ZF-R as a control. Binding of the activated RNA polymerase II was first evaluated by chromatin immunoprecipitation followed by qPCR (ChIP-qPCR) using an antibody targeting the phosphorylated form of the enzyme at serine 5 (RNAP II-S5P), which is normally associated with the initiation of RNA synthesis. Indeed, control cells showed a strong enrichment of RNAP II-S5P that peaked in proximity to the transcription start site (Figure 3A). In contrast, B2M-ZF-R expressing T cells exhibited a dramatic reduction of RNAP II-S5P signal near background levels, thus correlating with the strong MHC-I protein downregulation. Furthermore, as KRAB domains cooperate with chromatin-remodeling factors such as histone lysine-methyl transferases, we also interrogated the levels of histone 3 lysine 9 trimethylation (H3K9me3) on the *B2M* locus. Consistently, the loss of RNAP II-S5P in ZF-R-expressing T cells was accompanied with the acquisition of this repressive mark, whose enrichment also peaked around the TSS (Figure 3B). Overall, these data indicate that the silencing induced by the B2M ZF-R occurs through the epigenetic modulation of the local chromatin.

**Figure 3.**
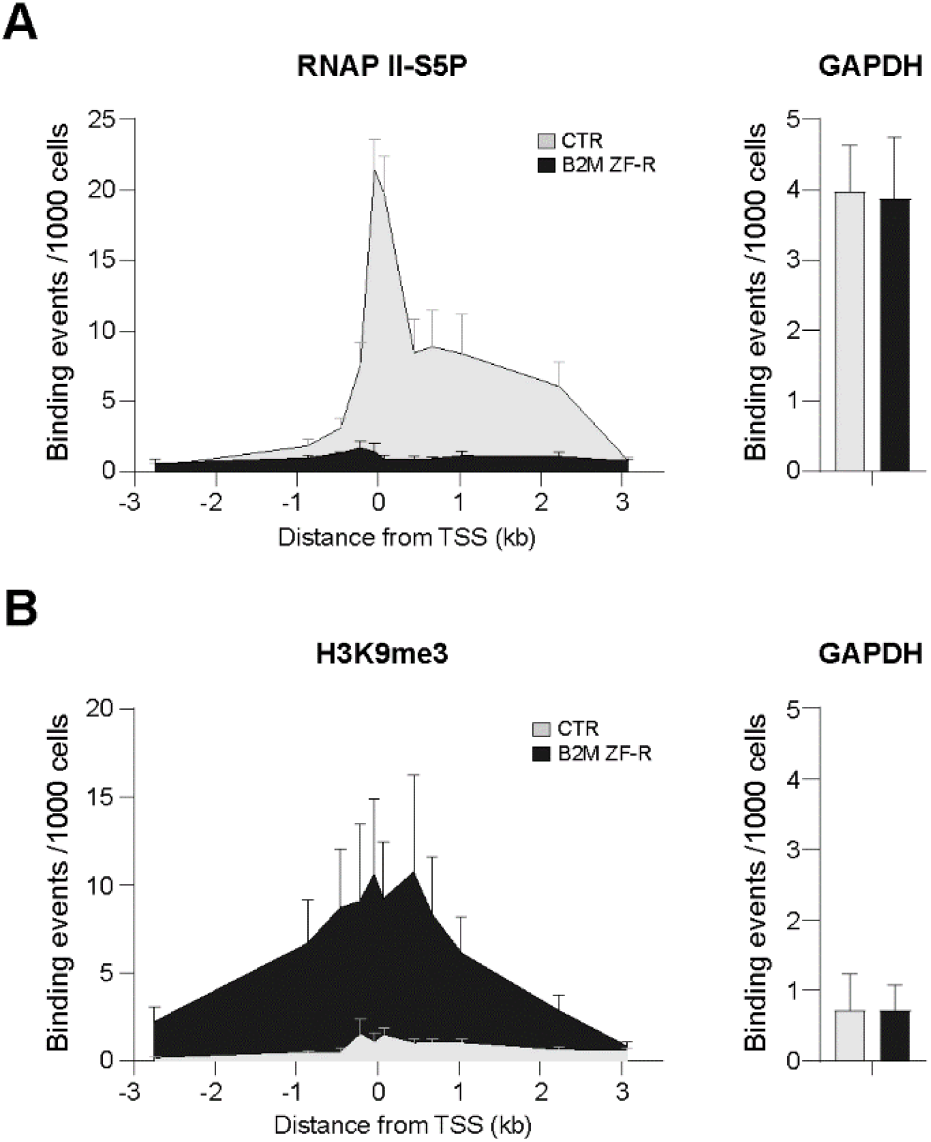
Epigenetic repression at the *B2M* and *CD5* targeted locus. Chromatin immunoprecipitation followed by qPCR (ChIP-qPCR) analysis for the phosphorylated form of RNA Pol II at serine 5 (RNAP II-S5P, A) and H3K9me3 epigenetic mark (B) performed on the *B2M* gene of control and B2M ZF-R-expressing T cells and on the unrelated *GAPDH* gene (Right histograms). Data show enrichment events per 1000 cells (n=3 independent T cell donors). Data are represented as mean ± SEM.

### B2M and CD5 co-repression efficiency and durability in ZF-R-expressing T cells

To monitor the durability of the ZF-R-mediated epigenetic repression, ZF-R-expressing T cells were injected in an immunodeficient mice model and monitored for ten weeks *in vivo* (Figure 4A). ZFR-expressing T cells were enriched based on dNGFR expression (>93%) and expanded for two weeks *in vitro*, showing a stable MHC-I and CD5 repression (Figure S6A-B), and a stable CD4/CD8 T cell ratio (Figure S6C). Then, these MHC-I(-) and/or CD5(-) T cells were injected intraperitoneally into NXG mice and human T cell engraftment (hCD45+) was monitored every week in the mouse peripheral blood (Gating strategy Figure S7). As expected, without any human cytokine support like interleukin-2, engraftment slowly decreased over time for the control and B2M ZFR-expressing T cells (Figure 4B). On the contrary, T cells negative for CD5 were able to proliferate efficiently (Figure 4B), highlighting in this setting the negative role of CD5 on T cell activation/proliferation as described previously.^36–38^ Interestingly, MHC-I and/or CD5 downregulation was very stable during the entire experiment, independently of the proliferation status (Figure 4C). Importantly, ZF-R-expressing T cells isolated from the spleen or the murine blood (Week 10) displayed the same repression profile as before injection (Figure 4D). These data suggest that the lentiviral vector is an optimal delivery system for sustained expression of multiplexed ZF-Rs, allowing durable and concomitant repression of multiple genes.

**Figure 4.**
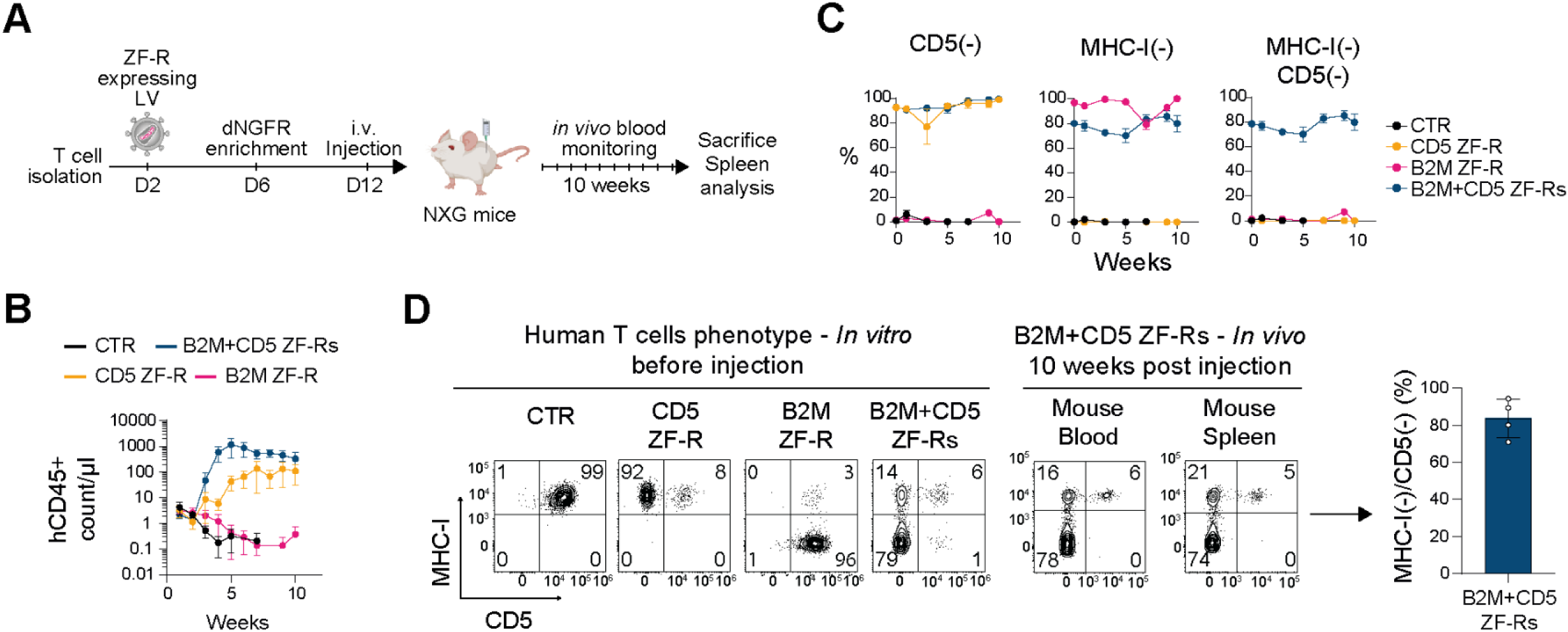
Efficient and durable MHC-I and CD5 repression in human ZF-R expressing T cells engrafted *in vivo*. A) Schematic of the *in vivo* experimental protocol. B) Monitoring of the engraftment efficiency of hCD45+ cells during 10 weeks. C) Durability of MHC-I and/or CD5 downregulation in ZF-R-expressing T cells injected in NXG mice (n=5/group). D) Representative FACS plots showing MHC-I and CD5 downregulation in dNGFR+ T cells before injection and 10 weeks post-injection in blood and spleen. On the right panel, the histogram represents the percentage of concomitant MHC-I(-) / CD5(-) T cells in spleen (n=5 mice). Data are represented as mean ± SEM.

### ZF-R-mediated downregulation of MHC-I and MHC-II in CAR-Tregs

To extend the evaluation of the ZF-R technology to different T cell subsets, ZF-Rs were delivered in CAR-expressing regulatory T cells (CAR-Tregs).^39^ Tregs were transduced with a vector expressing a CAR directed against the human HLA.A2 protein^40^ and two ZF-Rs directed against B2M and CIITA to downregulate MHC-I and MHC-II expression, respectively (Figure S2), an approach previously described to produce off-the-shelf hypoimmunogenic CAR-T cells.^24, 25^ As shown in Figure 5A-B, concomitant repression of B2M and CIITA genes led to the generation of around 70% of CAR-Tregs with no cell surface expression of MHC-I and MHC-II molecules. Constitutive expression of these ZF-Rs had no impact on CAR-Treg activation (*i.e.* CD69 expression), neither after stimulation through the TCR/CD3 complex using anti-CD3/CD28 beads nor through the CAR using a fluorescent dextramer coated with HLA.A2 proteins (Figure 5C). Furthermore, the phenotype of ZF-R-expressing CAR-Tregs remained stable *in vitro* (Figure 5D), strongly suggesting that the integrity and functionality of primary cells is not impacted by the constitutive expression of ZF-Rs *per se*.

**Figure 5.**
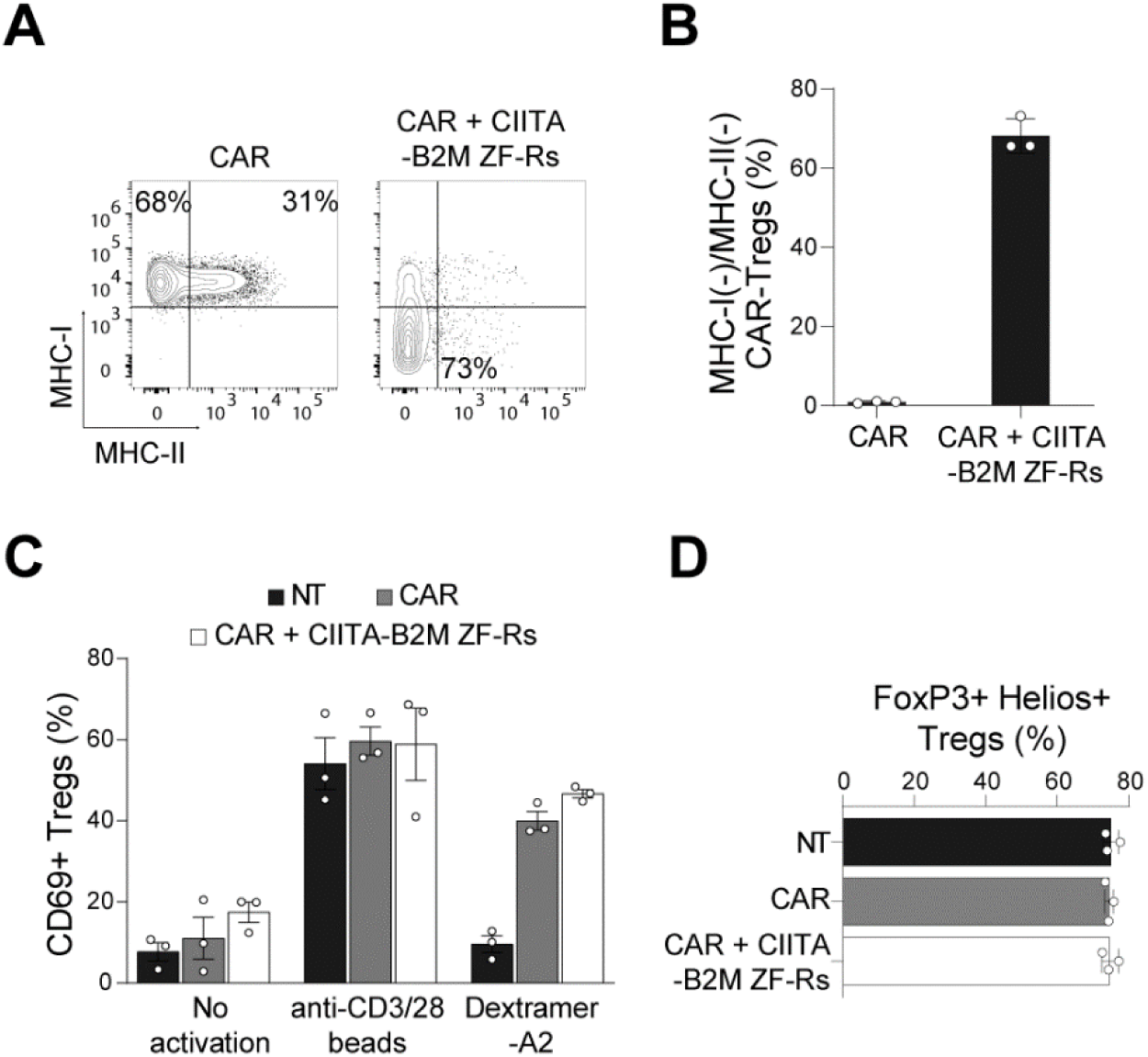
ZF-R-mediated downregulation of MHC-I and MHC-II in CAR-Tregs. A) Representative FACS plots of CAR-Tregs transduced with the CAR alone (CAR) or multiplexed with B2M and CIITA ZF-Rs showing co-repression of MHC-I and MHC-II expression at D7. B) Percentage of double negative MHC-I and MHC-II cells measured at D12. C) Expression of the CD69 activation marker was assessed by flow cytometry one day after incubation with either anti-CD3/CD28 beads to stimulate the TCR (positive control) or with Dextramer-A2 to stimulate the HLA-A2 CAR. D) The expression of FoxP3 and Helios Treg markers was assessed via intracellular staining after cell surface staining and gating on CD4+, CD25+ and CD127low positive cells. Histograms represent the mean ± SEM. Alle the data were obtained from 3 independent T cell donors.

### Assessment of the ZF-repressor specificity

Despite high efficiency and favorable safety profile of the research grade ZF-Rs tested in this study (i.e. no impact on cell viability, expansion, phenotype, and functionality), non-specific interactions of ZF-Rs to genomic DNA, described as off-target sites, can occur. The development of clinical grade ZF-Rs will require high on-target specificity and nearly undetectable off-targets. We have previously shown that an arginine residue in each zinc-finger DNA binding domain makes a non-specific contact with the DNA phosphate backbone; the mutation of this arginine to glutamine, known as phosphate-contact variant (PCV), can modulate DNA binding affinity and reduce the off-target binding of zinc-finger nucleases.^41^ To evaluate and improve the specificity profile of the research-grade B2M ZF-R, PCV variants of the parental B2M ZF-R were generated. First, a microarray analysis was performed to identify the number of deregulated genes (DEGs) in T cells transduced with the B2M-ZF-R LV compared to the empty LV vector control. As shown in Figure 6A, the microarray results confirmed the strong repression of the *B2M* gene and a limited number of other DEGs. The genomic sequence of the more significant DEGs were aligned with the on-target sequence, either fully (18 consecutive base pairs) or partially. Only the *PMVK* gene presented a significant sequence homology to the *B2M* on-target sequence (Figure 6B), with twelve consecutive base pairs, covering the first (ZF-1 and -2) and the second two-finger module (ZF-3 and -4). Therefore, one PCV mutation was incorporated in either ZF-2 (PCV2) or ZF-4 (PCV4) to decrease the affinity of one of the two modules implicated in this off-target. As shown in Figure 6C, microarray analysis of T cells expressing the PCV2 variant showed a strong improvement in the specificity profile and these results were confirmed by RT-qPCR (Figure S8). The PCV4 variant was excluded because its specificity profile was still harboring two DEGs. Of note, only one DEG persisted in presence of the PCV2 variant, namely *PATL2*, a gene located only 200bp upstream of the *B2M* locus (see Figure S9). RT-qPCR analysis showed a partial repression of PATL2 (around 65%). This deregulation of the *PATL2* gene could be the consequence of an on-target effect of the B2M ZF-R since the localization of the H3K9Me3 epigenetic mark is extended +/- ∼2 kb of the target site, as shown in Figure 3B. Finally, despite a decrease in the non-specific affinity of PCV2 and PCV4 for the genomic DNA, the on-target interaction and B2M repression was still highly efficient, leading to more than >90% of MHC-1 downregulation (Figure 6D).

**Figure 6.**
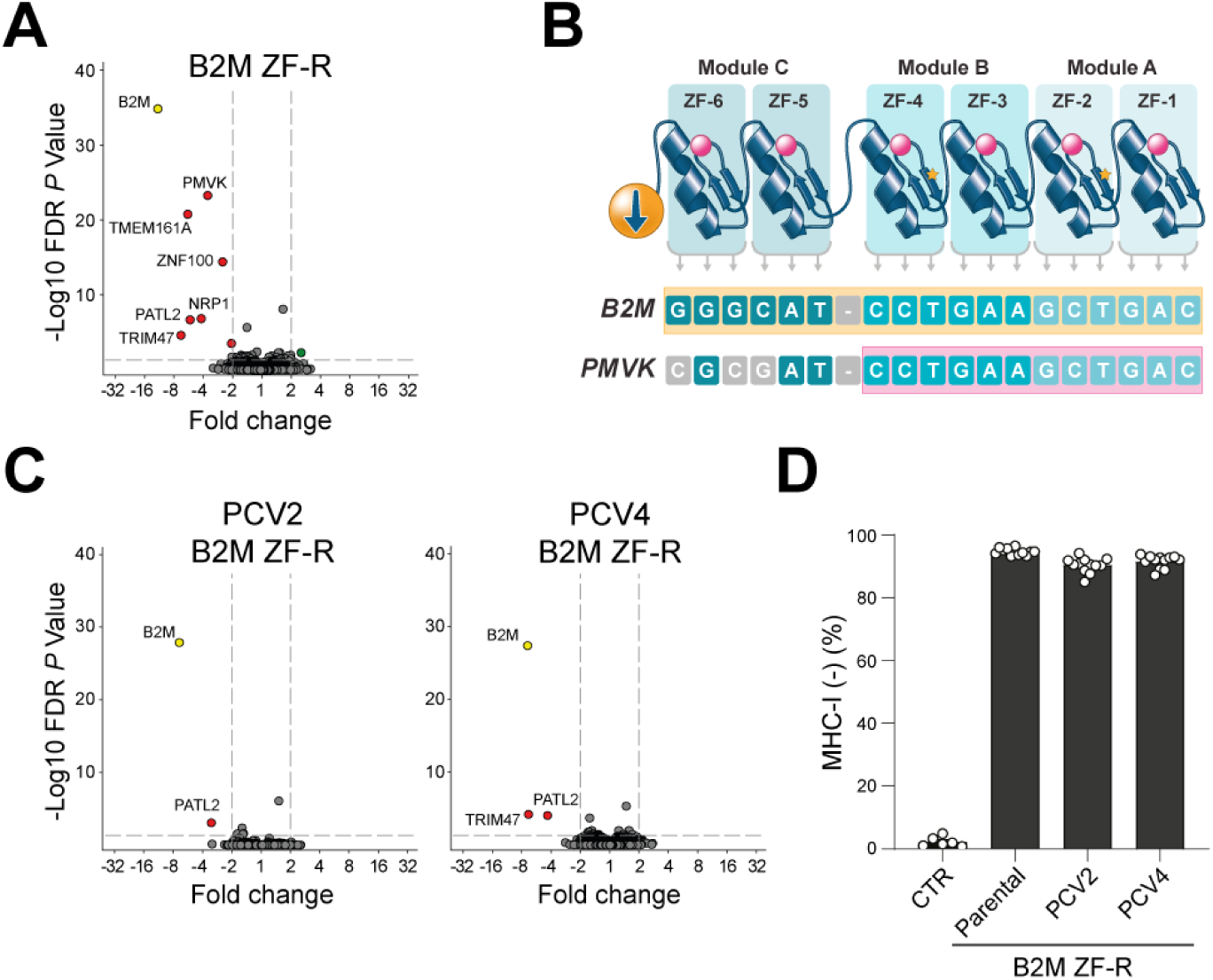
Design of phosphate-contact variants of B2M ZF-R to improve specificity. A) Volcano plot representation of pooled RNA array data from dNGFR-enriched T cells (n= 3 donors) transduced with parental B2M ZF-R (n=12) compared to empty LV vector (n=6). Threshold for significant deregulation (dotted lines) at false discovery rate (FDR)–adjusted *P* values < 0.05. The *B2M* target gene is marked with a yellow dot, other downregulated (red dots) or upregulated genes (green dots) more than >2 fold are annotated. B) Schematic of the sequence homology between B2M target DNA sequence and the deregulated PMVK gene. Zinc-finger proteins with Arg-to-Gln PCV mutation are highlighted with a yellow star. C) Volcano plot representation as in A with PCV2 and PCV4 B2M-ZF-R variants (n=12) compared to empty LV vector (n=6). D) Efficiency of MHC-I downregulation in dNGFR-enriched T cells in quadruplicate (n= 3 donors) expressing parental, PCV2 and PCV4 B2M ZF-R.

## DISCUSSION

In this study, we sought to design a versatile engineering strategy allowing concomitant and stable silencing of multiple genes in the same target cell. The development of engineered human-derived ZF-Rs for the epigenetic repression of target genes has the advantage of mirroring endogenous ZFP-KRABs already operating on the human genome. Furthermore, ZF-Rs lack the genotoxicity associated to nuclease-mediated gene silencing (i.e DNA double-strand breaks, translocations, etc.) and extend the silencing potential to noncoding RNAs. Interestingly, the number of highly active ZF-Rs targeting the TSS region of a given gene is variable, as exemplified here with B2M and CD5 ZF-R screenings. High chromatin accessibility for multiple CD5 ZF-Rs can be the consequence of low nucleosome occupancy, or high nucleosome occupancy with low stability and/or an increase in inter-nucleosomal spacing.^42–44^

The intracellular co-delivery of multiple epigenetic silencers for therapeutic applications can be achieved with HIV-1-derived LVs. They are well adapted for *ex vivo* engineering of hematopoietic cells, allowing permanent expression of silencers. Deployment of a single lentiviral particle to control multi-gene expression gives major advantages including: i) large cargo capacity, ii) concomitant delivery of all epigenetic repressors in each transduced cell, and iii) more simplicity in the process development and consequently a cheaper manufacturing cost. Nonetheless, the cloning design for multiple transgenes into the proviral DNA remains a challenge. Here, we showed that the bidirectional architecture is well adapted to express up to three ZF-Rs. Importantly, the type and the orientation of the two promoters are critical. Generation of MHC-I and CD5 co-repressed T cells was improved from 20 to 80% just by swapping the dual promoter architecture from EF1a-PGK to PGK200-EF1a. The EF1a promoter, known to be stronger than PGK, allows a better expression of the ZF-Rs. But we also observed that the EF1a promoter in sense orientation was negatively impacted by the proximity of the antisense PGK promoter, a side effect strongly attenuated by shortening the PGK promoter from 500 to only 200bp (PGK200). Based on these data, we hypothesized that a competition for common transcription factors between PGK and EF1a may occur.

Next, we observed that the constitutive expression of ZF-Rs after lentiviral integration led to stable gene repression during the 2-weeks period of culture *in vitro*. To extend this monitoring, ZF-R-expressing T cells were injected into immunodeficient mice and showed again a strong capacity to silence target genes during the entire follow-up period of ten weeks. This silencing efficiency was not modulated by the cell proliferation status, suggesting that the epigenetic marks (e.g. H3K9Me3) are constantly deposited at each proliferation cycle. Indeed, during the ZF-R screening with transient mRNA expression, the gene repression was lost in proliferating T cells after few days of culture.

To extend our evaluation of the ZF-R multiplexing technology, we designed a next-generation CAR-Treg approach. Converting polyclonal Tregs into antigen specific Tregs by introducing a CAR gained significant attention as a potential treatment option for autoimmune diseases and solid organ rejection.^39, 40^ We engineered CAR-Tregs co-expressing ZF-Rs targeting *B2M* and *CIITA* genes, both required for cell surface expression of MHC-I and MHC-II respectively.^24, 25^ The ZF-R mediated repression for both was potent and stable. The constitutive expression of B2M and CIITA ZF-Rs *per se* and the consequent downregulation of MHC-I and MHC-II did not impact either the CAR functionality or the Treg phenotype, expanding the field of application of ZF-Rs to diverse hematopoietic cells.

Finally, we shared a strategy to optimize the specificity of research grade ZF-R to generate potential clinical-grade candidates. As an example, we showed that human T cell transduction with the parental B2M ZF-R led to some DEGs as shown in microarray analysis. These genes could be true off-targets or biologically co-deregulated genes, downstream of *B2M*. Going through gene ontology studies, we did not find any link between B2M expression and the identified DEGs. We also performed an *in-silico* analysis of the DEGs for any homology with the on-target sequence and found only one off-target candidate, namely the *PMVK* gene, harboring twelve consecutive base pairs fully homologous to the on-target site. Based on this homology, a PCV mutation was incorporated into the Zinc-Finger 2 (Module A) and/or Zinc-Finger 4 (Module B) of the parental ZF-R. While PCV2 and PCV4 maintained a strong repression of B2M expression, the double mutant PCV2-4 was way less efficient to repress B2M (around 30%) and was therefore excluded from the study (data not shown). The PCV4 B2M ZF-R was also excluded due to a remaining DEG (i.e. *TRIM47*), while The PCV2 B2M ZF-R mutant showed a dramatic drop for all DEGs in the microarray analysis and in RT-qPCR validation assays. Altogether, these data confirm that the disruption of the non-specific interaction of the Zinc-Finger to the DNA phosphate backbone is an elegant and attractive approach to avoid unspecific DNA binding of ZFPs.^41^ Interestingly, both the parental and the PCV2 B2M ZF-R resulted in some deregulation of the surrounding *PATL2* gene, located only 200 bp upstream of the *B2M* promoter (Figure S9). While *B2M* was strongly repressed (>90%), *PATL2* gene was partially repressed (65% as compared with control). Due to its close proximity to *B2M*, *PATL2* gene repression could potentially be an on-target effect of the B2M ZF-R. This slight deregulation of the closest neighboring gene is probably the consequence of an “Halo” effect of the KRAB domain, a phenomenon previously observed for the *PTMS* gene following the repression of the LAG3 gene located ∼1kb downstream.^45^ This is important to note that the modulation of expression of *PATL2* did not have any impact on T cell phenotype, viability, expansion, and B2M repression stability. This is consistent with the absence of a functional role of *PATL2* gene in the biology of human T cells. Nonetheless, it highlights the importance of monitoring the expression of surrounding genes during the development of clinical grade repressors, either ZFP-KRAB, Cas9-KRAB, or TALE-KRAB, to evaluate any potential halo effect of the KRAB domain along the few kilobases of the target gene locus.

In conclusion, we were able to concomitantly deliver multiple functional epigenetic repressors in a single target cell. Considering that ZFP binding domains can also be fused to compact transcriptional activation domains (TADs),^11, 46^ this cell engineering approach opens the way to innovative therapeutic applications requiring concomitant transgene expression (e.g. CAR, TCR) and modulation of expression of multiple genes with a performant and cost-effective single lentiviral transduction event.

## MATERIAL AND METHODS

### Human primary cell and cell line culture

For screening of ZF-Repressors, human T cells were obtained from fresh Leukopaks purchased from STEMCELL Technologies (Seattle, WA). T cell isolation was performed on the CliniMACS system using anti-CD4 and CD8 magnetic beads (Miltenyi Biotec, San Jose, CA). Next, T cells were suspended in CryoStor 10 (StemCell Technologies, Seattle, WA), aliquoted, and stored in liquid nitrogen. After thawing, T cells were cultured in complete X-vivo15 (Lonza, Hayward, CA), supplemented with 100 IU/ml IL-2 (Thermo Fisher Scientific Waltham, MA), 5% Human AB Serum (Valley Biomedical, Winchester, VA), and activated with anti-CD3/CD28 Dynabeads (Thermo Fisher Scientific, Waltham, MA) at a bead:cell ratio of 1:3. For other experiments, primary cells were isolated from buffy coats of healthy donors (EFS, Marseille, France). Peripheral blood mononuclear cells (PBMCs) were isolated by Ficoll gradient centrifugation. T cells and Tregs were isolated by immunomagnetic selection. Briefly, T cells were isolated using the EasySep^TM^ Human T cell isolation kit (StemCell Tech., Saint-Egrève, France) and cultured in complete X-Vivo15 medium. Tregs were isolated using the EasySep human CD4+CD127lowCD25+ Regulatory T Cell Isolation Kit (StemCell Tech.) and cultured in X-Vivo15, supplemented with 1000 IU/mL IL-2, anti-CD3/CD28 dynabeads (1:1 ratio) and 100nM Rapamycin (Sigma-Aldrich, Saint-Quentin-Fallavier, France). Jurkat E6.1 cells (Sigma-Aldrich) were cultured in X-Vivo15 medium.

### ZF-Repressor design and screening

#### Design and assembly of ZF-Repressors

ZFP backbones, based on the human ZFP Zif268/EGR, were designed using an archive of prevalidated one- and two-finger modules as previously described.^47^ The resulting ZFPs were cloned into the pVAX vector (Thermo Fisher Scientific) to produce a fusion with KRAB repressor domains described in this study.

#### ZF-Repressor mRNA production and transfection

Templates for *in vitro* transcription were generated by PCR from pVAX-ZFP-KRAB plasmids (Fwd primer: GCAGAGCTCTCTGGCTAACTAGAG; Rev primer: T(60)CTGGCAACTAGAAGGCACAG). mRNA transcripts were synthesized using the mMESSAGE mMACHINE T7 ULTRA Transcription Kit (Thermo Fisher Scientific) as per the manufacturer’s instructions and purified using the Agencourt RNAClean XP Kit (Beckman Coulter, Brea, CA) on a Kingfisher instrument (Thermo Fisher Scientific). T cells were cultured in complete X-Vivo15 medium for three days prior to mRNA transfection. ZF-Repressor mRNA transfection was performed using 10ug of ZFP mRNA per 2×10E5 cells on a BTX Multi-well Electroporation 96-well plate with a 2mm well gap with a BTX ECM 830 Electroporator with Plate Handler HT-200 (BTX Harvard Apparatus, Holliston, MA). The transfection percentage was >90% as determined from co-transfection with a control GFP encoding mRNA. Forty-eight hours after transfection, cells were lysed and analyzed for gene expression by RT-qPCR.

#### Gene expression analysis using RT-qPCR

mRNA samples from ZF-R-transfected T cells were reverse transcribed to cDNA using a Power SYBR Green Cells-To-CT kit (Thermo Fisher Scientific) and qPCR reactions were performed using QuantiFast Multiplex PCR Master Mix (w/o ROX) (Qiagen, Redwood City, CA). Each biological replicate was assessed in technical quadruplicate on a 384-well CFX real-time qPCR instrument (Bio-Rad, Hercules CA) and analyzed using the Bio-Rad CFX Manager 3.1 software. The housekeeping genes EIF4A2 and ATP5B were used for RT-qPCR normalization. The RT-qPCR probe/primer sets used are listed in supplemental Table S1.

### Lentiviral vector production and titration

Lentiviral vectors were produced using the classical 4-plasmid lentiviral system. Briefly, HEK293T cells (Lenti-X, Ozyme, France) were transfected with plasmids expressing HIV-1 Gag/pol (pMDLg/pRRE), HIV-1 Rev (pRSV.Rev), the VSV-G glycoprotein (pMD2.G) (Didier Trono, EPFL, Switzerland), and a third-generation transfer plasmid expressing the transgene. 24 hrs post-transfection, viral supernatants were harvested, concentrated by centrifugation, aliquoted and frozen at -80°C for long term storage. Infectious titers expressed in transducing units per milliliter (TU/ml) were obtained after transduction of Jurkat T cells with a serial dilution of viral supernatants and transduction efficiency evaluated after 3-4 days by monitoring cell surface dNGFR expression using an anti-CD271-APC antibody (Miltenyi Biotec., Paris, France) or HLA-A*02 Dextramer-APC (Immudex, Copenhagen, Denmark) for HLA.A2 CAR.

### Lentiviral transduction and dNGFR enrichment

Lentiviral constructs used in this study are described in Figure S2. Lentiviral transduction was performed two days after T cell or Treg isolation and activation. Briefly, cell suspension (2×10^6^ cells/ml) were incubated with LV supernatants at a final concentration of 4×10^7^ TU/ml in complete X-Vivo15 medium. After 6 hours at 37°C and 5% CO2, cell suspensions were diluted eight times with fresh complete medium. After 3 to 5 days, transduction efficiencies were measured using flow cytometry to determine the percentage of dNGFR or CAR positive cells. For dNGFR enrichment, transduced cells were isolated with the EasySep Human CD271 Positive Selection Kit II with EasyEights™ EasySep™ Magnets (StemCell Technologies) according to the manufacturer protocol.

### In vitro Treg activation assay

In the activation assay, Tregs were stimulated for 24 hrs through the TCR (positive control) with anti-CD3/CD28-coated dynabeads (1 bead:2 cells ratio) (ThermoFisher Sci.) or through the HLA.A2-CAR with 2.5µL HLA-A2*02 dextramer (Immudex). Cells were then harvested and analyzed by flow cytometry.

### Vector copy number and mRNA quantification in LV-transduced T cells

The vector copy number (VCN) per cell was measured by qPCR. Genomic DNA (gDNA) was extracted using the DNeasy Blood and tissue kit (Qiagen, Hilden, Germany) according to manufacturer’s instructions. To determine the VCN, amplification of the human albumin (*ALB*) gene and the HIV-1 Psi vector sequence were performed using respectively custom qPCR HEX assay and custom qPCR FAM assay (Bio-Rad, Marnes-la-Coquette, France) with previously published primer sequences.^48^ qPCR reactions were performed using 200 ng of gDNA and the Master Mix Taqman Universal II (Applied Biosystems, Villebon-sur-Yvette, France) according to manufacturer’s recommendations. All reactions were performed using the CFX Opus 96 Dx System (Bio-Rad). A standard curve was used to convert Ct values to copy numbers by amplifying serial dilutions of a unique plasmid containing equimolar ratios of the ALB and PSI target sequences.

Expression levels of indicated mRNA transcripts in LV-transduced cells were analyzed by RT-qPCR. Total RNA was extracted from dNGFR-enriched cells using the RNAqueous total RNA isolation kit (Ambion, Austin, United States) and subsequently treated with Turbo DNA free-kit (Ambion). Reverse transcription was performed with Verso cDNA Synthesis kit (ThermoFisher Sci.). Gene-specific primers and probes are described in the Table S1 and S2. Experiments were carried out in duplicate for each data point. The relative quantification in gene expression was determined using the 2-ΔΔCt method.^49^ Using this method, fold changes in gene expression were normalized to the housekeeping GAPDH mRNA.

### Chromatin immunoprecipitation – qPCR assay

Chromatin immunoprecipitation followed by real-time quantitative PCR (ChIP-qPCR) was performed using the ChIP-IT PBMC kit (53042; Active Motif, Carlsbad CA) according to manufacturer’s instructions. Briefly, NGFR-enriched T cells from three different healthy donors (purity > 90%) were harvested and fixed with 1% formaldehyde for 10 min, lysed and prepared for sonication. Genomic DNA was fragmented to a mean length of 300–1500 bp using the Epishear probe sonicator (Active Motif) and was controlled on 1.5% agarose gel. Next, immunoprecipitation with 12 µg of fragmented chromatin was performed using the following antibodies from Abcam (Cambridge, UK): anti-H3 (ab1791), anti-RNA polymerase II CTD repeat YSPTSPS phospho S5 (ab5131); anti-H3K9me3 (ab8898) and an unrelated IgG control (ab171870). qPCR reactions were performed in duplicate to detect specific genomic regions using SYBR Green (BioRad) and specific primers (Table S3). Results were analyzed with the ChIP-IT qPCR Analysis kit (53029; Active Motif) to calculate binding events detected per 1000 cells. Pol II phospho S5 and H3K9me3 signals were normalized with H3 signals performed with anti-H3 antibody on the same chromatin.

### NXG mouse model for *in vivo* monitoring of human ZF-R-expressing T cells

NOD-Prkdc^scid^-IL2rg^Tm1^/Rj (NXG) mice were obtained from Janvier Labs (Le Genest-Saint-Isle, France). 8-week-old NXG mice were housed in containment isolators and habituated for one week prior to experimental use. The animal protocol was approved by the French animal ethics committee. Mice were randomly assigned to groups. For *in vivo* experiments, T cells were transduced at D2, dNGFR-enriched at D6, and injected intraperitoneally at D12 (3 ×10^6^ T cells per mouse). Throughout experiments, the body weight, and the graft-versus-host disease (GvHD) score were monitored twice a week by operators blinded to treatment. Once a week, blood samples were collected by retro-orbital sampling under local anesthesia. At the end of the experiment, mice were euthanized, tissue and blood samples were collected, and processed for immunophenotyping.

### Human primary cells immunophenotyping

For cellular immunophenotyping, T cells were stained with conjugated monoclonal antibodies (mAb) targeting CD3, CD4, CD8, CD5, CD271/NGFR (Miltenyi Biotec) MHC-I and MHC-II (BD Biosciences, Le Pont de Claix, France). For HLA-A2 CAR-Treg labeling, cells were stained at cell surface with conjugated mAb targeting CD4, CD25 and CD127 (Miltenyi Biotec) and the CAR was detected after incubation with a conjugated HLA-A2*02 dextramer (Immudex). Following fixation/permeabilization, Tregs were stained with conjugated mAb targeting the intracellular Helios (eBiosciences, Life technologies corp., Carlsbad, CA) and FoxP3 proteins (BD Biosciences). For *in vivo* experiments, mouse spleen, and bone marrow samples were passed through a 70µm cell strainer to obtain a single cell suspension. Red blood cells were lysed with red blood cell lysis buffer (Merck, Fontenay-Sous-Bois, France). Next, cells were incubated with mouse Fc block (BD Biosciences) and stained with the indicated antibodies. Marker expressions were measured by flow cytometry and analyzed using FlowJo software (BD Biosciences).

### Microarray studies

T cells were transduced on day 2 with the B2M-ZF-R LV or the empty LV control vector. Four days later, ZF-R-expressing T cells were enriched using the dNGFR marker and total RNA extracted on day 12. Total RNA was analyzed for gene expression changes using a microarray analysis (Clariom S Human HT, Thermo Fisher Scientific). Each replicate (100ng of total RNA) was processed according to the manufacturer’s protocols for sample preparation, hybridization, fluidics, and scanning. Robust multi-array average (RMA) was used to normalize the raw signal from each probe set. Fold change analysis was performed using Transcriptome Analysis Console 4.0 (Thermo Fisher Scientific) with “analysis type – expression (gene)” and “summarization –SST-RMA” options selected. Gene expression levels in ZF-R-expressing T cells were always compared to T cells transduced with an empty control LV. Change calls are reported for transcripts (probe sets) with a >2-fold difference in mean signal relative to control and an FDR P-value <0.05 [one-way analysis of variance (ANOVA) analysis and unpaired t test for each probe set]. *n* = 19,669 transcript clusters assessed.

### Statistics

Statistical analyses were performed using GraphPad Prism v.10.1.0 (GraphPad Software Inc., San Diego, CA, USA).

## DATA AVAILABILITY STATEMENT

All requests for data will be reviewed by Sangamo Therapeutics, Inc. to verify whether the request is subject to any intellectual property or confidentiality obligations. If deemed necessary, a material transfer agreement between the requestor and Sangamo Therapeutics, Inc. may be required for the sharing of some data. Any data that can be freely shared will be released.

## ACKNOWLEDGMENTS

The authors would like to thank the members of the animal care facility at Sangamo Ther. France (Valbonne, France) and the design & technology team for the design and synthesis of ZFP encoding libraries at Sangamo Ther. (Richmond, CA, USA). We also thank Caroline Courme (SciComCube, Valbonne, France) for graphical assistance.

## AUTHOR CONTRIBUTIONS

D.M., M.D., S.K.T., Y.Z., I.M., C.J., G.S., C.F.D., A.E.M., L.T., J.E., C.N. and M.H. performed experiments. D.M., M.D., S.K.T., Y.Z., I.M., C.J., G.S., C.F.D., A.E.M., J.E., L.T., C.N., M.H., A.R. and D.F. designed experiments, analyzed and interpreted data. G.D.D., J.D.F. and M.d.l.R. contributed to the discussions. M.D., D.M., S.K.T., A.R. and D.F. wrote the manuscript. D.F. supervised the study.

## DECLARATION OF INTERESTS

All the authors are current or former Sangamo Therapeutics employees. Sangamo Therapeutics has filed a patent application covering the technology described in this paper.

## REFERENCES

1. Ecco, G, Imbeault, M, and Trono, D (2017). KRAB zinc finger proteins. Development 144: 2719–2729.

2. Senft, AD, and Macfarlan, TS (2021). Transposable elements shape the evolution of mammalian development. Nat Rev Genet 22: 691–711.

3. Yang, P, Wang, Y, and Macfarlan, TS (2017). The Role of KRAB-ZFPs in Transposable Element Repression and Mammalian Evolution. Trends Genet 33: 871–881.

4. Iyengar, S, and Farnham, PJ (2011). KAP1 protein: an enigmatic master regulator of the genome. J Biol Chem 286: 26267–26276.

5. Zhang, Y, Sun, Z, Jia, J, Du, T, Zhang, N, Tang, Y, et al. (2021). Overview of Histone Modification. Adv Exp Med Biol 1283: 1–16.

6. Gersbach, CA, and Perez-Pinera, P (2014). Activating human genes with zinc finger proteins, transcription activator-like effectors and CRISPR/Cas9 for gene therapy and regenerative medicine. Expert Opin Ther Targets 18: 835–839.

7. Alerasool, N, Segal, D, Lee, H, and Taipale, M (2020). An efficient KRAB domain for CRISPRi applications in human cells. Nat Methods 17: 1093–1096.

8. Thakore, PI, D’Ippolito, AM, Song, L, Safi, A, Shivakumar, NK, Kabadi, AM, et al. (2015). Highly specific epigenome editing by CRISPR-Cas9 repressors for silencing of distal regulatory elements. Nat Methods 12: 1143–1149.

9. Cong, L, Zhou, R, Kuo, YC, Cunniff, M, and Zhang, F (2012). Comprehensive interrogation of natural TALE DNA-binding modules and transcriptional repressor domains. Nat Commun 3: 968.

10. Zhang, Z, Wu, E, Qian, Z, and Wu, WS (2014). A multicolor panel of TALE-KRAB based transcriptional repressor vectors enabling knockdown of multiple gene targets. Sci Rep 4: 7338.

11. Gommans, WM, Haisma, HJ, and Rots, MG (2005). Engineering zinc finger protein transcription factors: the therapeutic relevance of switching endogenous gene expression on or off at command. J Mol Biol 354: 507–519.

12. Jamieson, AC, Miller, JC, and Pabo, CO (2003). Drug discovery with engineered zinc-finger proteins. Nat Rev Drug Discov 2: 361–368.

13. Rebar, EJ, and Pabo, CO (1994). Zinc finger phage: affinity selection of fingers with new DNA-binding specificities. Science 263: 671–673.

14. Paschon, DE, Lussier, S, Wangzor, T, Xia, DF, Li, PW, Hinkley, SJ, et al. (2019). Diversifying the structure of zinc finger nucleases for high-precision genome editing. Nat Commun 10: 1133.

15. Charlesworth, CT, Deshpande, PS, Dever, DP, Camarena, J, Lemgart, VT, Cromer, MK, et al. (2019). Identification of preexisting adaptive immunity to Cas9 proteins in humans. Nat Med 25: 249–254.

16. Mehta, A, and Merkel, OM (2020). Immunogenicity of Cas9 Protein. J Pharm Sci 109: 62–67.

17. Simhadri, VL, McGill, J, McMahon, S, Wang, J, Jiang, H, and Sauna, ZE (2018). Prevalence of Pre-existing Antibodies to CRISPR-Associated Nuclease Cas9 in the USA Population. Mol Ther Methods Clin Dev 10: 105–112.

18. Garriga-Canut, M, Agustin-Pavon, C, Herrmann, F, Sanchez, A, Dierssen, M, Fillat, C, et al. (2012). Synthetic zinc finger repressors reduce mutant huntingtin expression in the brain of R6/2 mice. Proc Natl Acad Sci U S A 109: E3136–3145.

19. Scott, TA, Soemardy, C, Ray, RM, and Morris, KV (2023). Targeted zinc-finger repressors to the oncogenic HBZ gene inhibit adult T-cell leukemia (ATL) proliferation. NAR Cancer 5: zcac046.

20. Wegmann, S, DeVos, SL, Zeitler, B, Marlen, K, Bennett, RE, Perez-Rando, M, et al. (2021). Persistent repression of tau in the brain using engineered zinc finger protein transcription factors. Sci Adv 7.

21. Zeitler, B, Froelich, S, Marlen, K, Shivak, DA, Yu, Q, Li, D, et al. (2019). Allele-selective transcriptional repression of mutant HTT for the treatment of Huntington’s disease. Nat Med 25: 1131–1142.

22. Labbe, RP, Vessillier, S, and Rafiq, QA (2021). Lentiviral Vectors for T Cell Engineering: Clinical Applications, Bioprocessing and Future Perspectives. Viruses 13.

23. Poletti, V, and Mavilio, F (2021). Designing Lentiviral Vectors for Gene Therapy of Genetic Diseases. Viruses 13.

24. Depil, S, Duchateau, P, Grupp, SA, Mufti, G, and Poirot, L (2020). ’Off-the-shelf’ allogeneic CAR T cells: development and challenges. Nat Rev Drug Discov 19: 185–199.

25. Hu, X, Manner, K, DeJesus, R, White, K, Gattis, C, Ngo, P, et al. (2023). Hypoimmune anti-CD19 chimeric antigen receptor T cells provide lasting tumor control in fully immunocompetent allogeneic humanized mice. Nat Commun 14: 2020.

26. Geisler, C, Kuhlmann, J, and Rubin, B (1989). Assembly, intracellular processing, and expression at the cell surface of the human alpha beta T cell receptor/CD3 complex. Function of the CD3-zeta chain. J Immunol 143: 4069–4077.

27. D’Oro, U, Munitic, I, Chacko, G, Karpova, T, McNally, J, and Ashwell, JD (2002). Regulation of constitutive TCR internalization by the zeta-chain. J Immunol 169: 6269–6278.

28. Dai, Z, Mu, W, Zhao, Y, Jia, X, Liu, J, Wei, Q, et al. (2021). The rational development of CD5-targeting biepitopic CARs with fully human heavy-chain-only antigen recognition domains. Mol Ther 29: 2707–2722.

29. Cameron, P, Coons, MM, Klompe, SE, Lied, AM, Smith, SC, Vidal, B, et al. (2019). Harnessing type I CRISPR-Cas systems for genome engineering in human cells. Nat Biotechnol 37: 1471–1477.

30. Shaimardanova, AA, Kitaeva, KV, Abdrakhmanova, II, Chernov, VM, Rutland, CS, Rizvanov, AA, et al. (2019). Production and Application of Multicistronic Constructs for Various Human Disease Therapies. Pharmaceutics 11.

31. Szymczak, AL, and Vignali, DA (2005). Development of 2A peptide-based strategies in the design of multicistronic vectors. Expert Opin Biol Ther 5: 627–638.

32. Liu, Z, Chen, O, Wall, JBJ, Zheng, M, Zhou, Y, Wang, L, et al. (2017). Systematic comparison of 2A peptides for cloning multi-genes in a polycistronic vector. Sci Rep 7: 2193.

33. Amendola, M, Venneri, MA, Biffi, A, Vigna, E, and Naldini, L (2005). Coordinate dual-gene transgenesis by lentiviral vectors carrying synthetic bidirectional promoters. Nat Biotechnol 23: 108–116.

34. Golding, MC, and Mann, MR (2011). A bidirectional promoter architecture enhances lentiviral transgenesis in embryonic and extraembryonic stem cells. Gene Ther 18: 817–826.

35. Frigault, MJ, Lee, J, Basil, MC, Carpenito, C, Motohashi, S, Scholler, J, et al. (2015). Identification of chimeric antigen receptors that mediate constitutive or inducible proliferation of T cells. Cancer Immunol Res 3: 356–367.

36. Alotaibi, F, Rytelewski, M, Figueredo, R, Zareardalan, R, Zhang, M, Ferguson, PJ, et al. (2020). CD5 blockade enhances ex vivo CD8(+) T cell activation and tumour cell cytotoxicity. Eur J Immunol 50: 695–704.

37. Dasu, T, Qualls, JE, Tuna, H, Raman, C, Cohen, DA, and Bondada, S (2008). CD5 plays an inhibitory role in the suppressive function of murine CD4(+) CD25(+) T(reg) cells. Immunol Lett 119: 103–113.

38. Tarakhovsky, A, Kanner, SB, Hombach, J, Ledbetter, JA, Muller, W, Killeen, N, et al. (1995). A role for CD5 in TCR-mediated signal transduction and thymocyte selection. Science 269: 535–537.

39. Requejo Cier, CJ, Valentini, N, and Lamarche, C (2023). Unlocking the potential of Tregs: innovations in CAR technology. Front Mol Biosci 10: 1267762.

40. Proics, E, David, M, Mojibian, M, Speck, M, Lounnas-Mourey, N, Govehovitch, A, et al. (2023). Preclinical assessment of antigen-specific chimeric antigen receptor regulatory T cells for use in solid organ transplantation. Gene Ther 30: 309–322.

41. Miller, JC, Patil, DP, Xia, DF, Paine, CB, Fauser, F, Richards, HW, et al. (2019). Enhancing gene editing specificity by attenuating DNA cleavage kinetics. Nat Biotechnol 37: 945–952.

42. Mansisidor, AR, and Risca, VI (2022). Chromatin accessibility: methods, mechanisms, and biological insights. Nucleus 13: 236–276.

43. Mieczkowski, J, Cook, A, Bowman, SK, Mueller, B, Alver, BH, Kundu, S, et al. (2016). MNase titration reveals differences between nucleosome occupancy and chromatin accessibility. Nat Commun 7: 11485.

44. Rando, OJ, and Chang, HY (2009). Genome-wide views of chromatin structure. Annu Rev Biochem 78: 245–271.

45. Wilken, MS, Ciarlo, C, Pearl, J, Schanzer, E, Liao, H, Biber, BV, et al. (2021). Quantitative dialing of gene expression via precision targeting of KRAB repressor. bioRxiv: 2020.2002.2019.956730.

46. Bhatt, B, Garcia-Diaz, P, and Foight, GW (2023). Synthetic transcription factor engineering for cell and gene therapy. Trends Biotechnol.

47. Urnov, FD, Rebar, EJ, Holmes, MC, Zhang, HS, and Gregory, PD (2010). Genome editing with engineered zinc finger nucleases. Nat Rev Genet 11: 636–646.

48. Hacein-Bey Abina, S, Gaspar, HB, Blondeau, J, Caccavelli, L, Charrier, S, Buckland, K, et al. (2015). Outcomes following gene therapy in patients with severe Wiskott-Aldrich syndrome. JAMA 313: 1550–1563.

49. Livak, KJ, and Schmittgen, TD (2001). Analysis of relative gene expression data using real-time quantitative PCR and the 2(-Delta Delta C(T)) Method. Methods 25: 402–408.

