## Supplementary Material for "Epigenetic control of multiple genes with a single lentiviral vector encoding transcriptional repressors fused to compact zinc finger arrays"

*D. Monteferrario, M. David et al.*

### Supplemental Tables

**Table S1. List of commercial assays used to monitor mRNA expression of the indicated genes.**

| Gene | Assay ID | Dye | Supplier |
|---|---|---|---|
| **Probes used to measure gene expression in mRNA electroporated T cells** | | | |
| *B2M* | Hs00187842_m1 | FAM | Thermo Fisher Sci. |
| *CD5* | Hs00204397_m1 | FAM | |
| *CD3z* | Hs00609515_m1 | FAM | |
| *CIITA* | Hs00172106_m1 | FAM | |
| *EIF4A2* | Hs.PT.58.3053665 | HEX | Integrated DNA Technologies |
| *ATP5B* | Hs.PT.58.20452786 | Cy5 | |
| **Probes used to measure gene expression in LV transduced T cells** | | | |
| *B2M* | qHsaCIP0029872 | FAM | Bio-Rad |
| *CD5* | qHsaCIP0029162 | FAM | |
| *GAPDH* | qHsaCEP0041396 | HEX | |
| *TRIM47* | qHsaCEP0025151 | FAM | |
| *TMEM161A* | qHsaCEP0040651 | FAM | |
| *ZNF100* | qHsaCEP0057965 | FAM | |
| *PATL2* | qHsaCIP0025914 | FAM | |
| *PMVK* | qHsaCIP0027910 | FAM | |

**Table S2. List of home-made primers and probes used to measure mRNA expression of the indicated genes.**

| Gene | Primer/ probe | Sequence | Dye | Supplier |
|---|---|---|---|---|
| *ZNF10 (KOX1)* | Fw | ACACTGGTGACCTTCAAGGA | FAM | Bio-Rad |
| | Rv | CTCCACCAGCCAGGGC | | |
| | Probe | TGGACTTCACCAGGGAGGAGTGGAAG | | |
| *NRP1* | Fw | ATACAGGTAGACTTGGGCCTTC | FAM | |
| | Rv | GTTGGAGCTAACGTCGATCTTG | | |
| | Probe | CACGGCTGTCGGGACACAGGGCGCC | | |

**Table S3. List of primers used for ChIP-qPCR.**

| Gene | Location (bp from TSS) | Primer sequence |
|---|---|---|
| *B2M* | -2749 | CCTGTTGATGTATGTTGGATCG |
| | | CCAAGGTATTTGCCCAAGAGAA |
| | -863 | AGTCTCTTAGCCTTTGTTTCCC |
| | | GGCTGGCACATAGTAGGTAC |
| | -463 | AGCTGTCCTCAGGATGCTT |
| | | ACGGAAACCCAGAAGATTAAGA |
| | -221 | GAAGTCCTAGAATGAGCGCC |
| | | ATGCACTAGACTGGGTGAGT |
| | -48 | CCTCTCTCTAACCTGGCACT |
| | | GAATGCTGTCAGCTTCAGGA |
| | 67 | CATTCGGGCCGAGATGTC |
| | | GGAGGGTAGGAGAGACTCAC |
| | 440 | GACAAAGTTTAGGGCGTCGA |
| | | ACGCGTTCACAAACCTCAG |
| | 662 | CGCTAAGTTCGCATGTCCTA |
| | | CATTCTACAAACGTCGCGTG |
| | 1025 | GAAGTCAAGCCTAACCAGGG |
| | | GGAGAATCTCACGCAGAAGG |
| | 2225 | CGCCTTCCCTCAAACAGAG |
| | | CCCATCAAGAGGTGGATTGG |
| | 3064 | GCAGGCTGATCACTTGAAGT |
| | | GACTACAGGCACACACCAC |
| *GAPDH* | -86 | GAAAGTAGGGCCCGGCTACTAG |
| | | CTGCGGGCTCAATTTATAGAAAC |

### Supplemental Figures

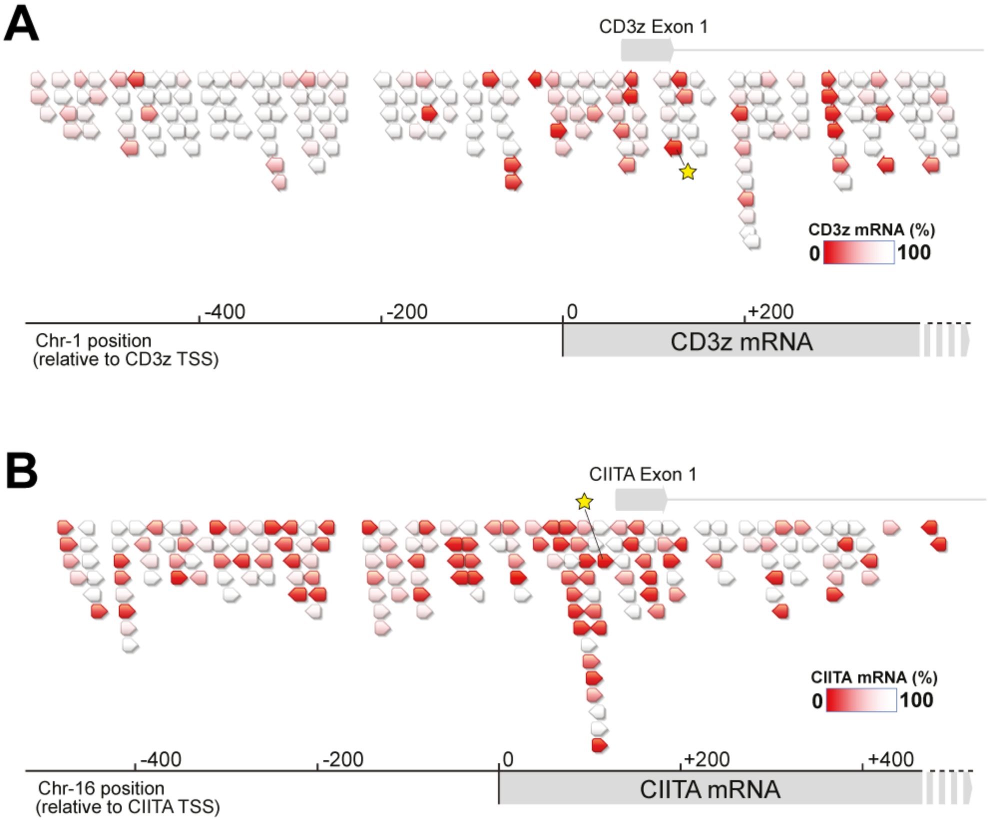


**Figure S1. Schematic of CD3zeta and CIITA ZF-Repressors screening.** Same schematic as Figure 1 representing the binding location and repression efficiency of ZF-Rs targeting the transcription start site (TSS) region of *CD3z* (A) and *CIITA* (B) genes in human T cells. The selected lead ZF-Rs are highlighted with a yellow star.

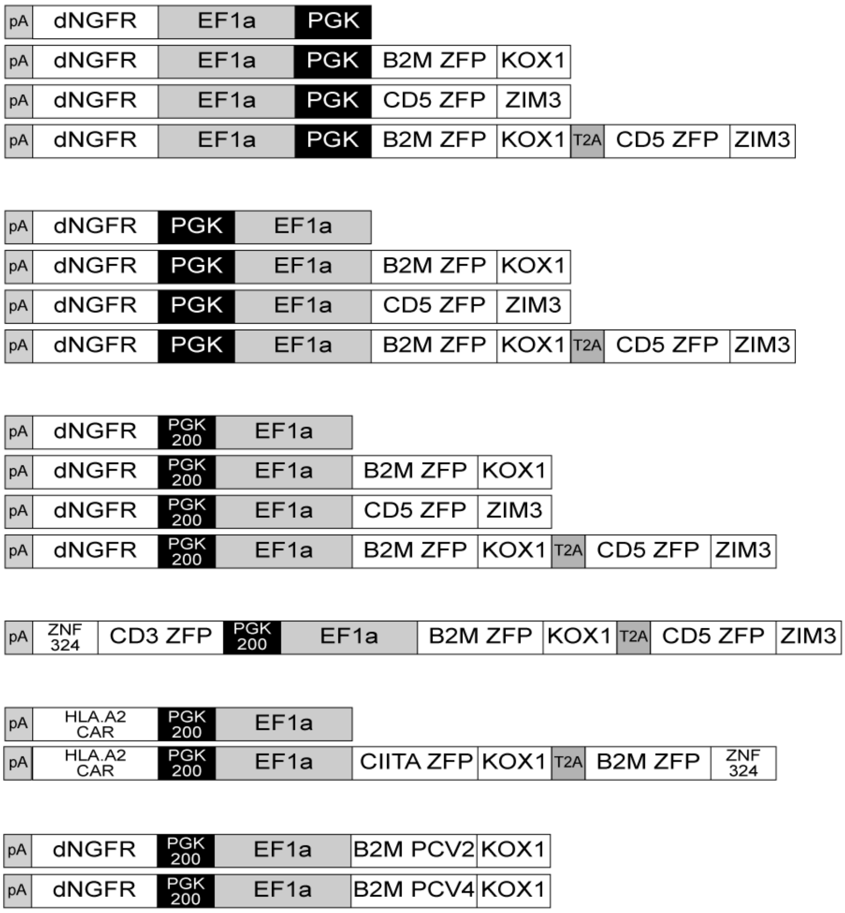


**Figure S2. Schematic representations of the lentiviral constructs used in this study.** pA, SV40 polyA; dNGFR, truncated human nerve growth factor receptor; PGK, human phosphoglycerate kinase promoter; PGK200, short 200bp PGK promoter; EF1a, human elongation factor 1 alpha promoter; HLA.A2 CAR, anti-human HLA.A2 chimeric antigen receptor. B2M zinc finger protein (ZFP) is fused either to the KOX1 or ZNF324 KRAB domain, CIITA ZFP is fused to KOX1, CD5 ZFP is fused to ZIM3 and CD3zeta ZFP is fused to ZNF324.

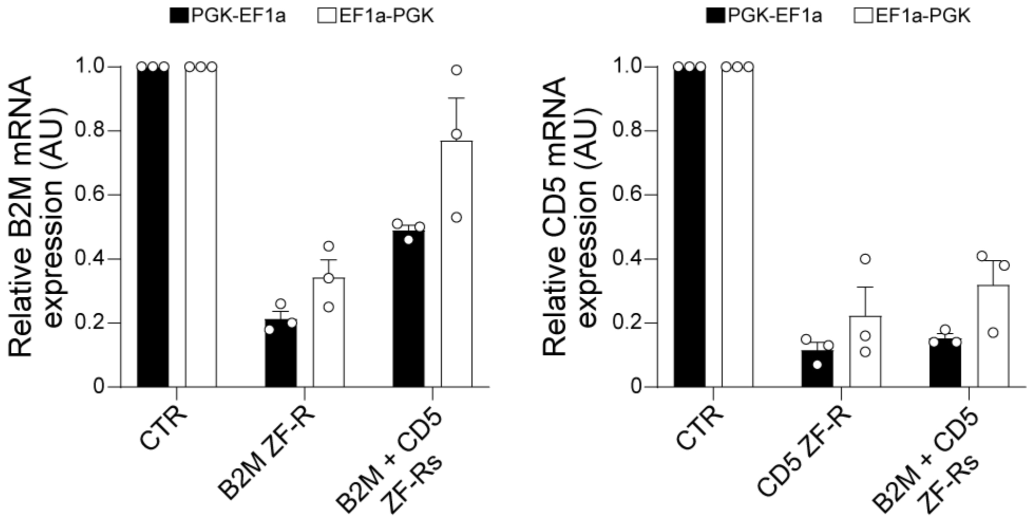


**Figure S3: Relative mRNA expression of *B2M* and *CD5* genes in ZF-R-expressing T cells.** B2M and CD5 ZF-R were cloned into the pLV.PGF-EF1a (black) or pLV.EF1a-PGK (white) lentiviral backbone in simplex or duplex format. Repression efficiency was assessed by measuring *B2M* (left panel) and *CD5* (right panel) mRNA transcripts expression (AU, arbitrary unit) in dNGFR-enriched cells at D10 post-transduction. Data are represented as mean ± SEM (n=3 donors, white circles).

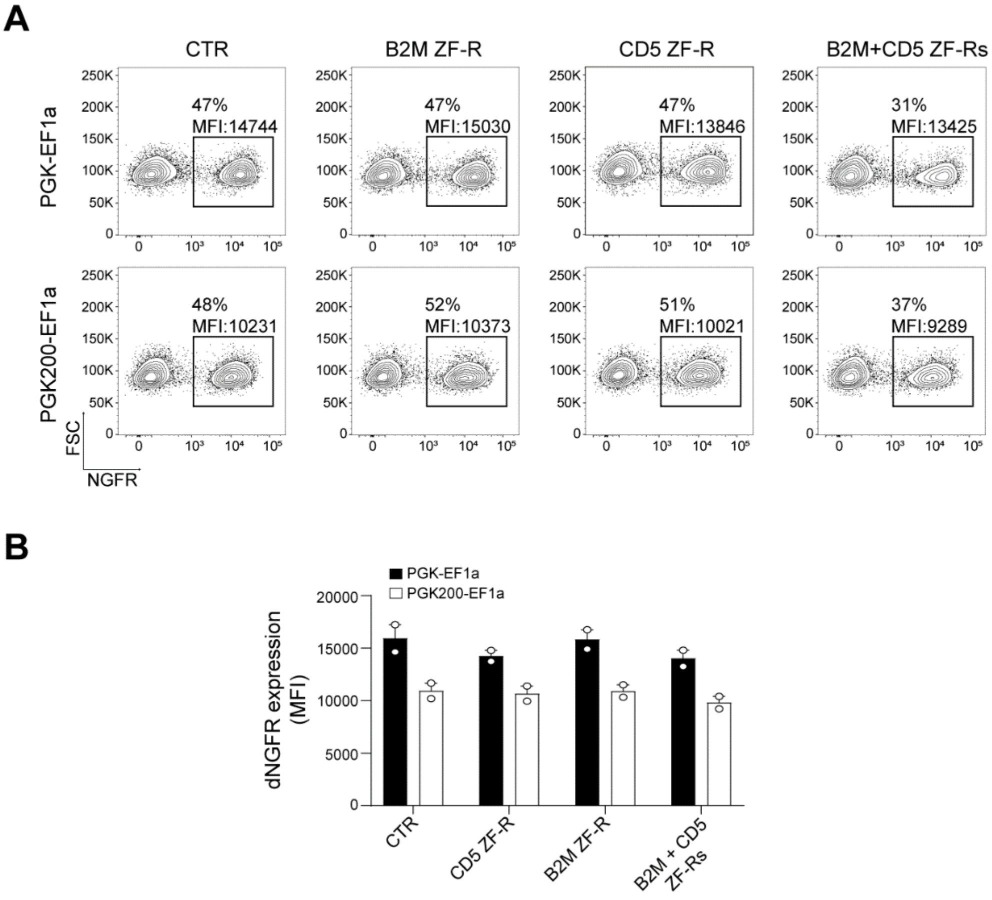


**Figure S4. No major difference in dNGFR expression under the control of the PGK versus PGK200 promoter.** Human T cells transduction with the indicated bidirectional LV (PGK-EF1a or PGK200-EF1a) expressing B2M, CD5 or B2M+CD5 ZF-Rs. dNGFR expression (MFI) was assessed by flow cytometry at D6 post-transduction. A) Representative FACS plots and B) histograms of dNGFR expression represented as the mean ± SEM (n=2 donors, white circles).

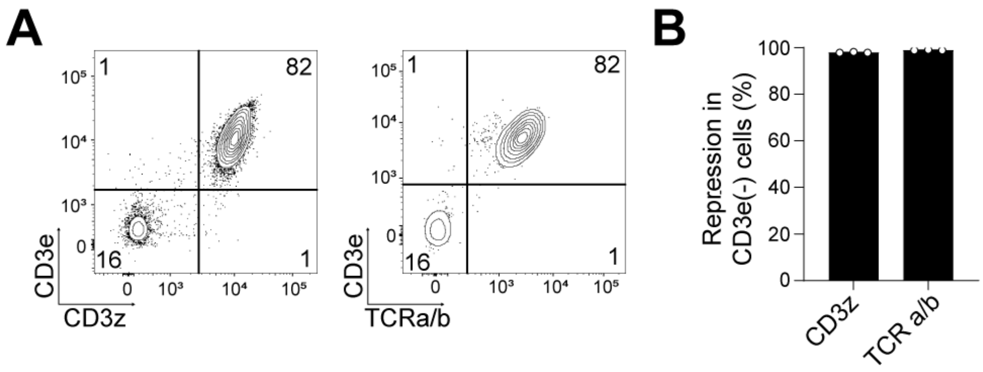


**Figure S5. Downregulation of TCR/CD3 complex in CD3z ZF-R-expressing T cells.**

A) Representative FACS plots of T cells transduced with CD3z ZF-R multiplexed with B2M and CD5 ZF-Rs showing that the repression of CD3z expression resulted in the simultaneous loss of CD3e and TCR a/b expression at cell surface. B) Quantification of CD3z and TCR a/b repression within the CD3e negative T cell population (n=3) represented in panel A.

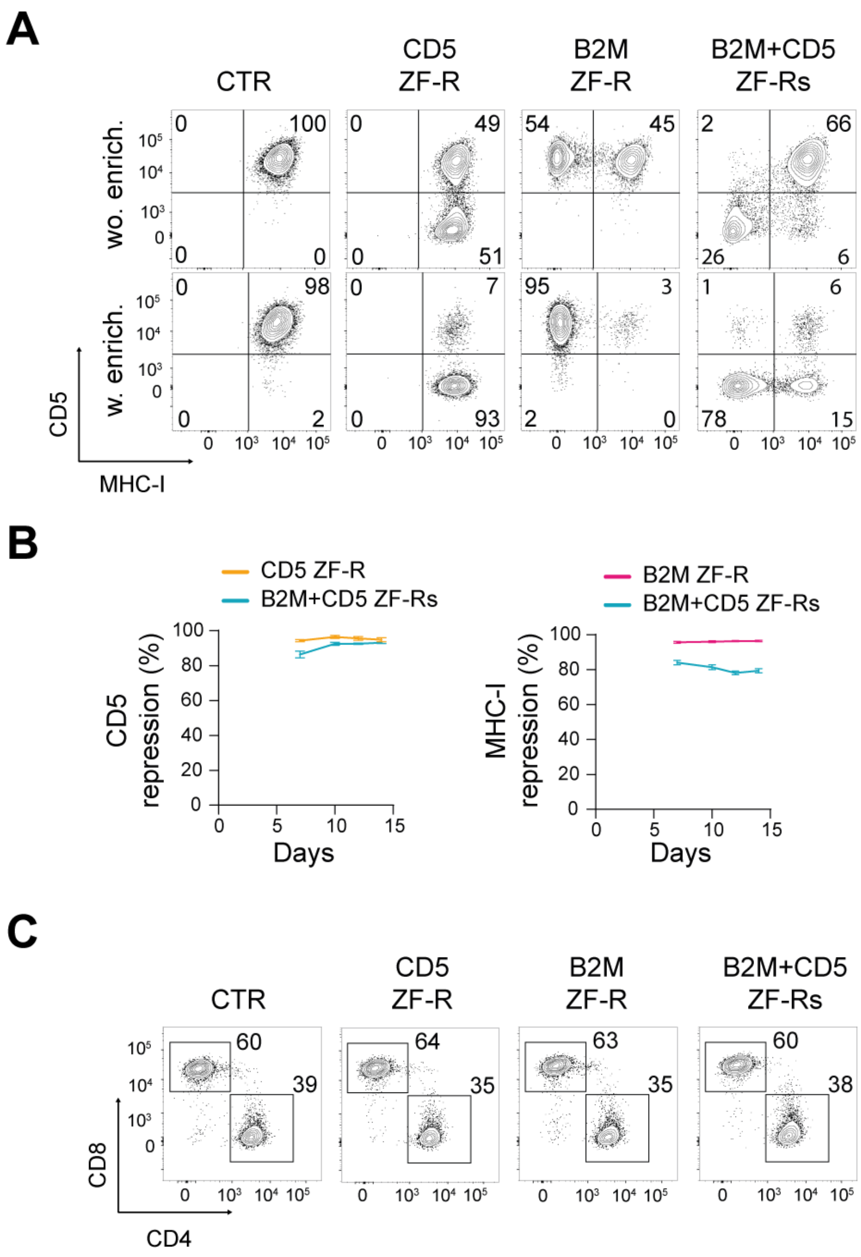


**Figure S6. Characterization of human ZF-R expressing T cells before *in vivo* injection.** Four days post-transduction with B2M, CD5 or B2M+CD5 ZF-R-expressing LV, dNGFR positive T cells were enriched via magnetic cell sorting. A) Comparison of MHC-I and CD5 expression levels at D12 with (w) or without (wo) dNGFR enrichment. B) Kinetics of MHC-I and CD5 repression efficiency in T cells after dNGFR enrichment at D4. Cells were cultured for 14 days with a reactivation step at D7 with anti-CD3/CD28 beads. C) Flow cytometry profile of CD4+ versus CD8+ T cells before *in vivo* injection at D12.

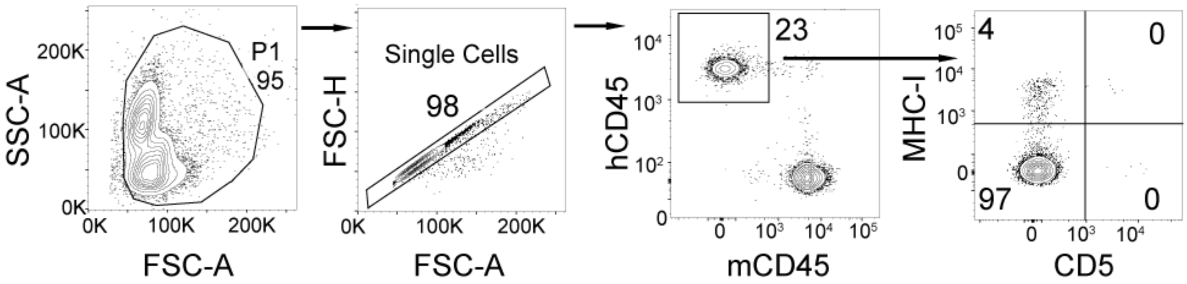


**Figure S7. Representative gating strategy to analyze *in vivo* engraftment of human T cells in blood.** After a gating strategy for single cells in forward-scatter area (FSC-A) versus height (FSC-H), human T cells were selected based on hCD45 expression. Next, repression efficiency was monitored after gating on the MHC-I(-) and/or CD5(-) population. The same strategy has been applied for human T cells monitored in murine spleen.

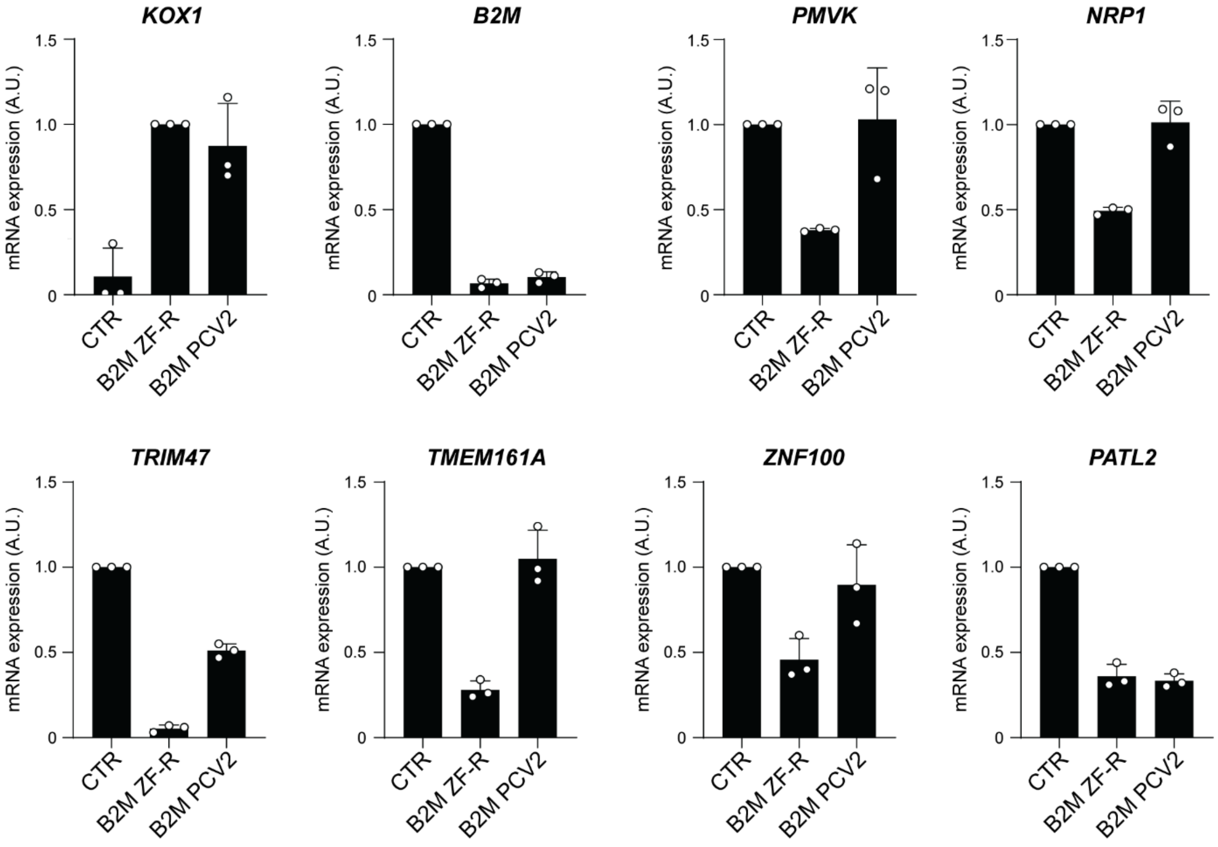


**Figure S8. Relative mRNA expression of DEGs in ZF-R-expressing T cells.** Relative quantification by RT-qPCR of mRNA transcript expression (AU, arbitrary unit) of *ZNF10*, *B2M*, *PMVK*, *NRP1*, *TRIM47*, *TMEM161A*, *ZNF100* and *PATL2* in dNGFR-enriched cells transduced with LV expressing no ZF-R (CTR), parental ZF-R (B2M ZF-R) or PCV2 B2M ZF-R (B2M PCV2). Data are represented as mean ± SEM (n=3 donors). For *ZNF10* (aka KOX1), the parental B2M ZF-R containing KOX1 sequence is used as the control and reflects the expression level of the ZF-R mRNA.

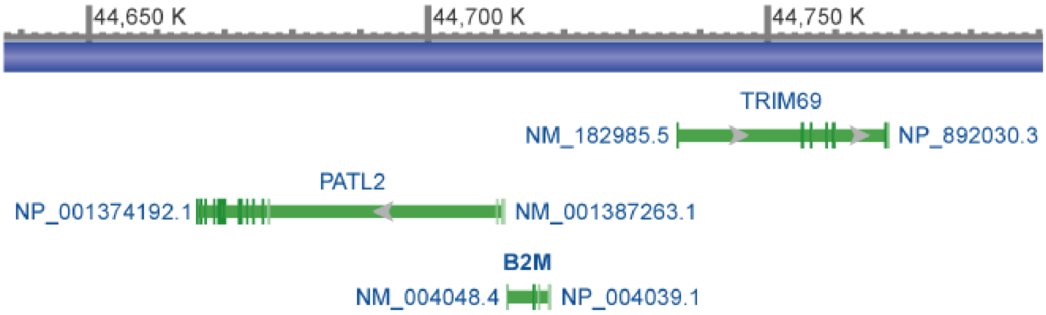


**Figure S9. Genomic region of the human *B2M* gene.** *Homo sapiens* genomic region encompassing the *PATL2*, *B2M* and *TRIM69* genes on chromosome 15. For each gene, NCBI references of the mRNA (NM) and protein (NP) are provided.

### Graphical Abstract

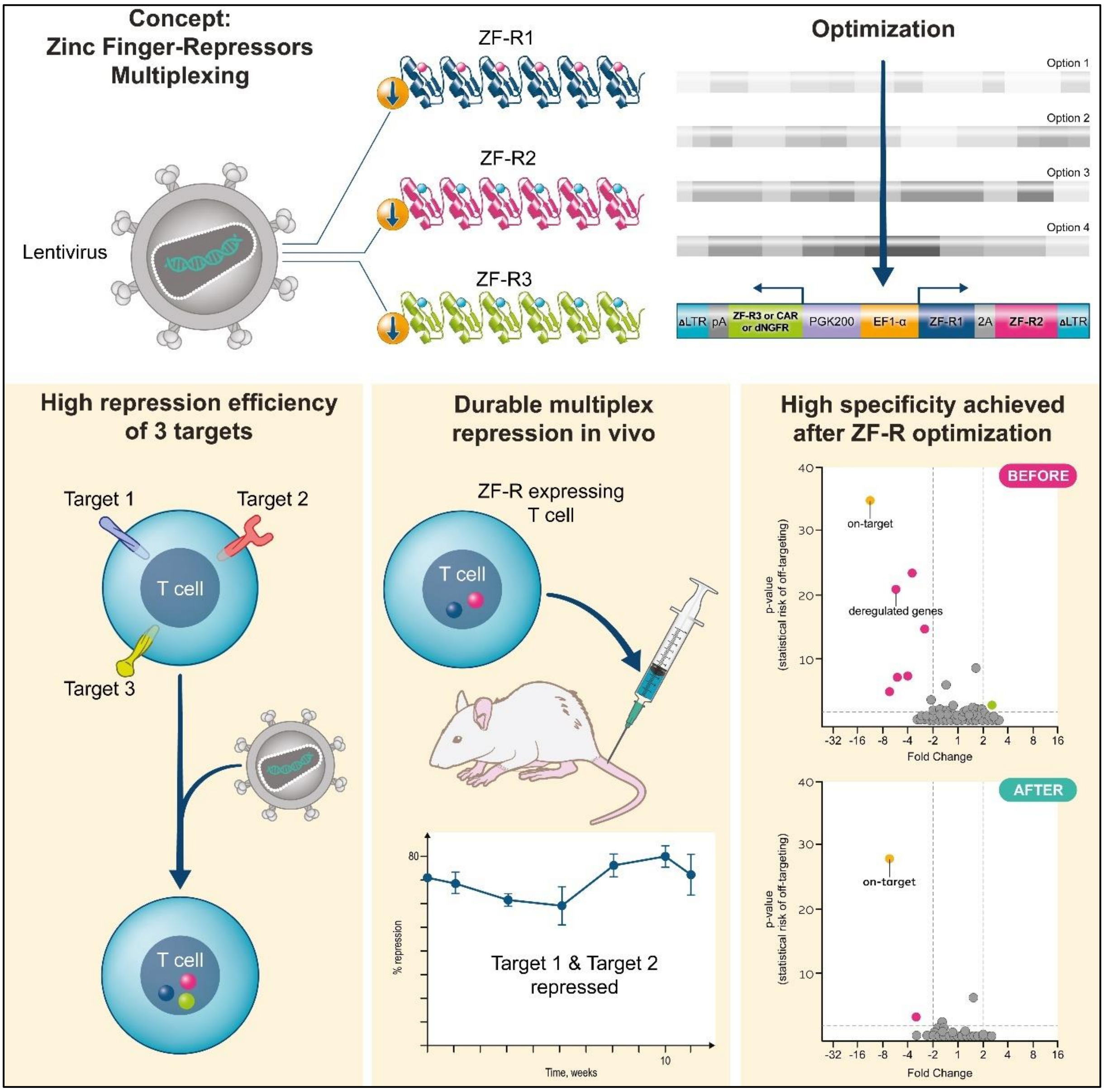
